# Could rare amino acids regulate enzymes abundance?

**DOI:** 10.1101/021295

**Authors:** Valentina Agoni

**Affiliations:** University of Pavia – Italy

## Abstract

The number of every enzyme is the result of many mechanisms of regulation, at transcriptional and translational levels. We asked if the presence of rare amino acids in the sequence of an enzyme could represent a limiting factor in the balance between enzymes synthesis and enzymes efficiency. To try to find a correlation we analyzed the amino acids sequences of different housekeeping and metabolic enzymes.

---

Like in economical model of pricing there is an equilibrium point between marginal cost and marginal revenue curves in the quantity vs price plot, similarly these concepts can be applied to enzymes synthesis and its energetic cost.

The number of copies of a certain enzyme is the result of many mechanisms of translational, transcriptional and post-translational modification regulation.

Gene expression can be regulated at several different levels, for example (1-17):

- Initiation of transcription by repressor or activator proteins
- Premature termination of transcription by attenuation
- Initiation of translation by an antisense RNA
- Chromatin domains
- Modification of DNA
- RNA transport
- mRNA degradation
- Post-translational by proteolysis or modification of the gene product

The frequency of the different amino acids in proteins (Figure 1) is known and these frequencies essentially reflect their frequency in nature and food, with the exception of arginine (18-20).

**Figure 1.**
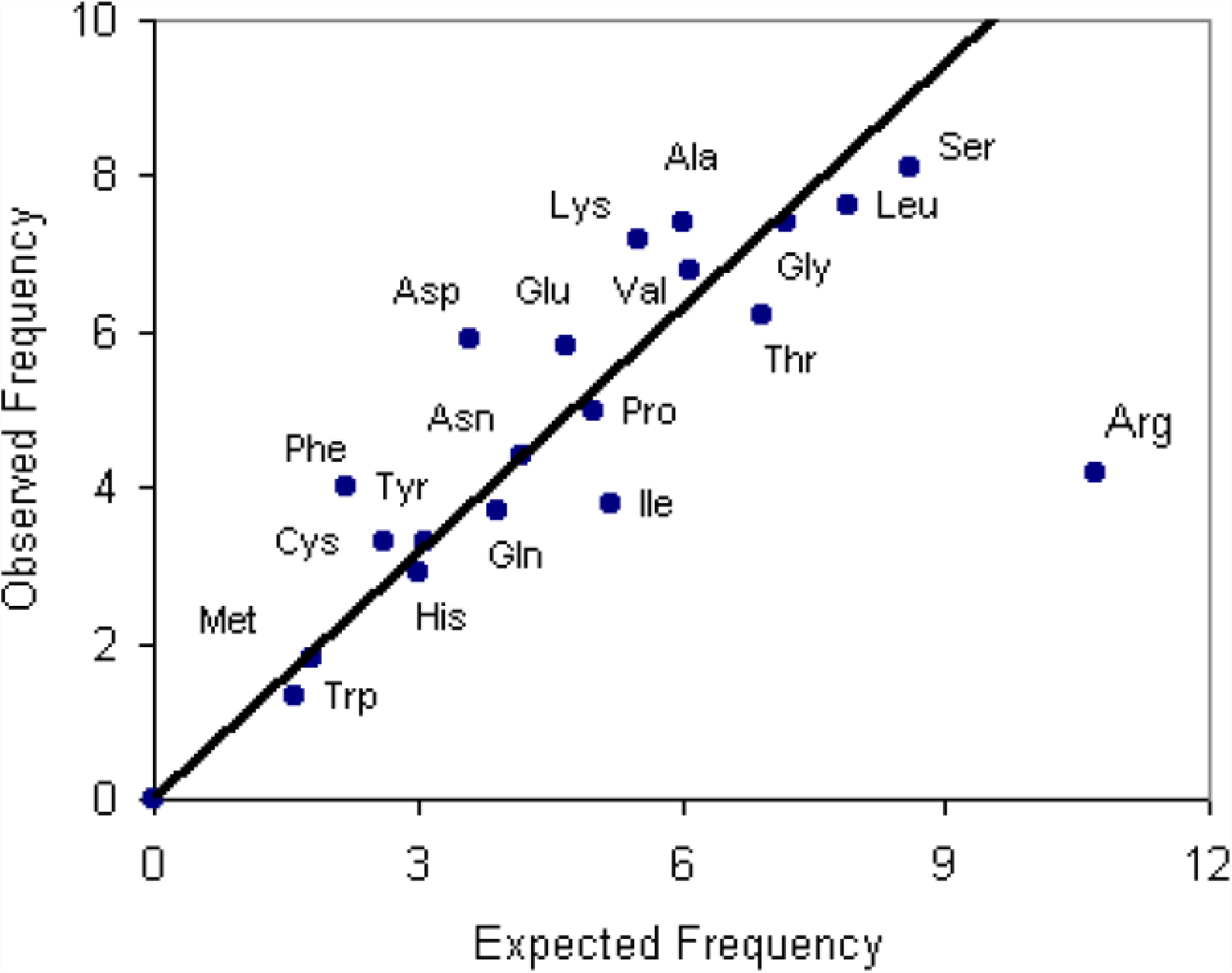
Amino acids expected frequency against the observed frequency in proteins (http://www.tiem.utk.edu/∼gross/bioed/webmodules/aminoacid.htm).

Essential amino acids in humans are:

Phe Val Thr Trp Met Leu Ile Lys His

and they correspond to the less used in protein sequences.

The amino acids frequencies are related to the proportion of alpha-helices and beta-strands in a protein.

In particular the amino acids more prone to create of alpha-helices are:

Met Ala Leu Glu Lys

While the more abundant in beta-strands are:

Tyr Phe Trp Thr Val Ile

We hypothesize that the frequency of each amino acid in proteins could represent a mechanism of regulation of enzymes relative abundance.

Here we consider the rarer and the more abundant amino acids: Trp (W) and Ser (S) respectively in housekeeping (h.k.) and metabolic or other enzymes.

H.k. enzymes vary their number of very little amount while the abundance of other enzymes can be much more variable reflecting physiological conditions (21-22).

According to our hypothesis the frequency of rare amino acids in h.k. enzymes should be higher respect to other enzymes.

For example Trp - the rarer amino acid in proteins-number should be lower in citrate synthase respect to GAPDH (glyceraldehyde-3-phosphate dehydrogenase).

However we can claim that there are no evidences of this correlation. The results are reported in the attachment of this article.

In other words, at least in principle, there is no relation between amino acids frequencies and the regulation of enzymes number.

